# Theoretical proposal for restoration of hand motor function utilizing plasticity of motor-cortical interhemispheric interaction

**DOI:** 10.1101/2024.05.30.596572

**Authors:** Hideki Nakano, Yandi Tang, Tomoyo Morita, Eiichi Naito

## Abstract

After stroke, the poorer recovery of motor function of upper extremities compared to other body parts is a longstanding problem. Based on our recent functional MRI evidence on healthy volunteers, this perspective paper proposes systematic hand motor rehabilitation utilizing the plasticity of interhemispheric interaction between motor cortices and following its developmental rule. We first discuss the effectiveness of proprioceptive intervention on the paralyzed (immobile) hand synchronized with voluntary movement of the intact hand to induce muscle activity in the paretic hand. In healthy participants, we show that this bilateral proprioceptive-motor coupling intervention activates the bilateral motor cortices (= bilaterally active mode), facilitates interhemispheric motor-cortical functional connectivity, and augments muscle activity of the passively-moved hand. Next, we propose training both hands to perform different movements, which would be effective for stroke patients who become able to manage to move the paretic hand. This bilaterally different movement training may guide the motor cortices into left-right independent mode to improve interhemispheric inhibition and hand dexterity, because we have shown in healthy older adults that this training reactivates motor-cortical interhemispheric inhibition (= left-right independent mode) declined with age, and can improve hand dexterity. Transition of both motor cortices from the bilaterally active mode to the left-right independent mode is a developmental rule of hand motor function and a common feature of motor function recovery after stroke. Hence, incorporating the brain’s inherent capacity for spontaneous recovery and adhering to developmental principles may be crucial considerations in designing effective rehabilitation strategies.

## 1. Introduction

The poorer recovery of motor function of upper extremities after stroke compared to other body parts is a longstanding problem (Lee et al., 2015). This perspective paper proposes systematic hand motor rehabilitation utilizing the plasticity of interhemispheric interaction between the motor cortices and following its developmental rule, based on recent neuroscientific evidence.

### 1.1 Motor-cortical bilaterally active mode is a spontaneous brain reaction after stroke

The motor cortex (precentral gyrus), which includes the primary motor cortex (M1) and dorsal premotor cortex (PMD), is the executive locus of motor control; damage to this area or descending tracts from this area can cause severe motor paralysis of the limbs. However, brain plasticity allows for functional recovery even in adult brains (Nudo, 2003; Grefkes and Fink, 2020). Interestingly, this functional recovery does not occur randomly. For example, when a patient manages to move a paretic hand after stroke, the contralesional motor cortex ipsilateral to the hand is often recruited in addition to the ipsilesional one, so that bilateral activity can be observed (Jang et al., 2004; Grefkes and Fink, 2020). This bilaterally active mode is a spontaneous brain reaction observed relatively early after stroke (within 10 days), and can be considered a state whereby the brain is searching for a new motor control pathway (including a pathway from the ipsilateral motor cortex) by trial and error to move the paretic hand. In other words, this bilateral mode is the first step toward restoring motor function (Grefkes and Fink, 2020). Recent animal studies have suggested that this bilaterally active mode is caused by disinhibition of interhemispheric inhibition between the left and right motor cortices (Yamaguchi et al., 2023) by acetylcholine modulating GABAergic interneurons (Handa et al., 2024).

### 1.2 Higher plasticity of motor-cortical interhemispheric inhibition

In the brain of typically developed young adults, there are interhemispheric facilitatory and inhibitory circuits between the motor cortices (Ni et al., 2020). When young adults perform simple unilateral movements (e.g. simple button pressing with a finger or simple hand alternating extension-flexion movement), the contralateral motor cortex is usually activated, and the ipsilateral motor cortex inhibited (Morita et al., 2019, 2021, 2023; Miura et al., 2024), probably due to interhemispheric inhibition between the two motor cortices (Mullinger et al., 2014). On the other hand, when young adults perform complex unilateral movements (e.g. stick-spinning or ball rotation with multiple fingers), there is activity in the ipsilateral motor cortex (especially in the PMD) in addition to the contralateral activity (Loibl et al., 2011; Uehara et al., 2012; Miura et al., 2024). Thus, the human brain adaptively controls movement by flexibly and plastically altering interhemispheric inhibition between the two motor cortices.

### 1.3 Developmental rule of motor-cortical interhemispheric inhibition and hyper-adaptation

The interhemispheric inhibition is not innate. During childhood, interhemispheric inhibition between the left and right motor cortices is still immature, maturing during adolescence (Ciechanski et al., 2017; Morita et al., 2019, 2021). Hence, the motor cortex before adolescence is in a bilaterally active mode, but this begins to change in adolescence to a left-right independent mode that allows the left and right hands to move independently. On the other hand, interhemispheric interaction between the two motor cortices can be greatly affected by training. We have recently shown that a top wheelchair racing Paralympian who trained from school age for special training in wheelchair racing, which requires bimanually synchronized upper-limb movements, shows a bilaterally active mode even in adulthood. She showed bilateral motor-cortical activations even during a simple alternating extension-flexion movement of the right hand, which should be called hyper-adaptation phenomenon rarely seen in typically-developed people (Morita et al., 2023). This finding inspired an intervention using bimanually synchronized movements that can act on the plasticity of interhemispheric interaction between the motor cortices.

In this paper, we first propose the effectiveness of an intervention to facilitate the motor cortices into bilaterally active mode, i.e., passive movement of one hand synchronized with voluntary movement of the other hand, based on our recent findings in healthy younger adults. Next, we introduce our previous findings on healthy older adults, and propose the effectiveness of training both hands to perform different movements to guide the motor cortices into left-right independent mode to improve interhemispheric inhibition and hand dexterity. By doing so, this paper provides theoretical and systematic framework for the interventions that utilize the higher plasticity of the motor-cortical interhemispheric interaction and that follow its developmental rule.

## 2 Induction of muscle activity utilizing bilaterally active mode

To move a paralyzed hand, it is first necessary to allow muscle activity in the paretic hand to emerge. Here, we consider an intervention that maximizes the bilaterally active mode, possibly the first step in the restoration of motor function after stroke. In the case of a hemiplegic patient who cannot move the right hand but can move the left hand, we propose a method in which his/her right hand is moved passively synchronized with voluntary movement of the left hand (Figure 1A).

**Figure 1.**
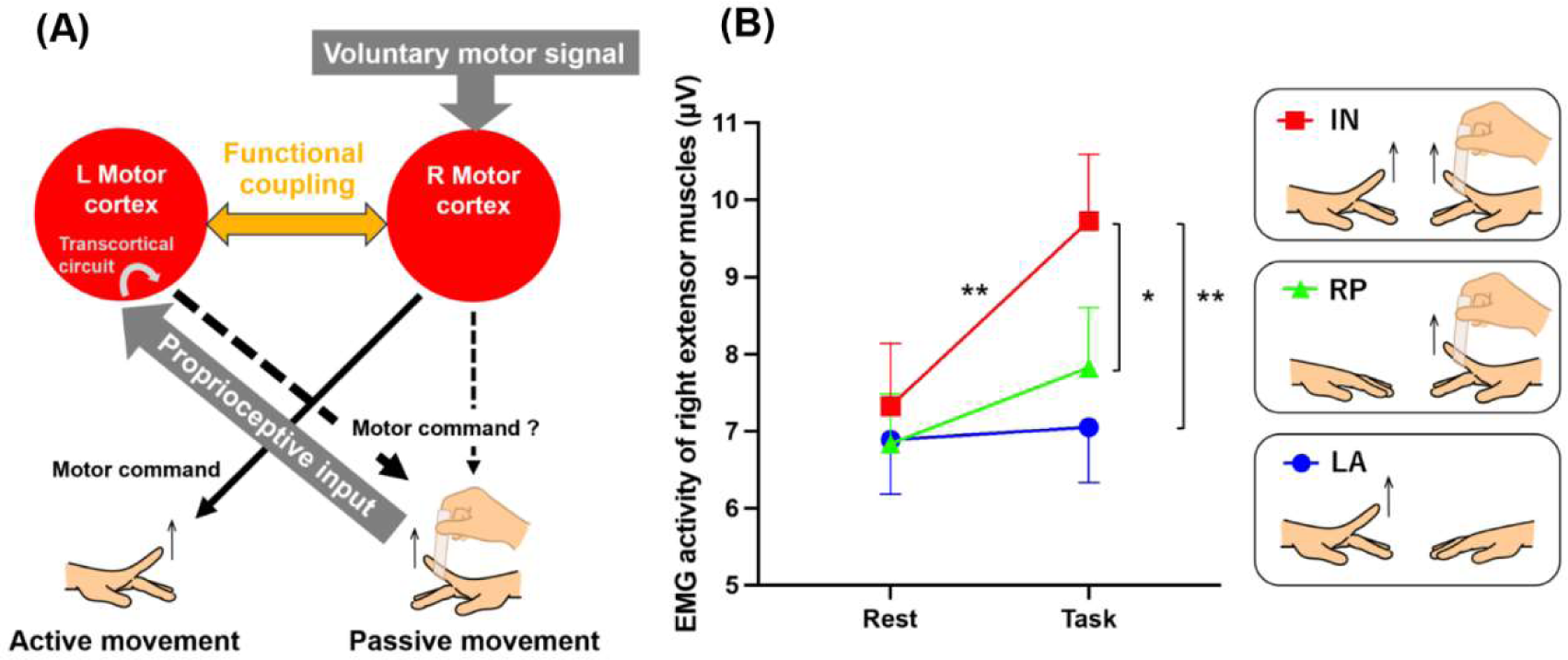
Possible neural mechanisms underlying passive extension of the right index finger synchronized with the voluntary extension of the left finger (A) and EMG results (B). (A) We hypothesize that passive movement of the right hand synchronized with voluntary movement of the left hand will lead the left and right motor cortices to a bilaterally active mode, and interhemispheric functional connectivity must be enhanced between the two motor cortices. This enhanced functional coupling may allow the bilateral motor cortices to share the voluntary motor signal input to the right motor cortex and the proprioceptive input to the left motor cortex, promoting sensory-motor associations between them. If the sensory-motor association occurs in the left motor cortex, the voluntary motor signal may effectively activate the intrinsic transcortical circuit between the left motor cortex and the right hand muscles, leading to muscle activity increase in the right passively-moved finger. If this association also occurs in the right motor cortex, the cortex might also become part of the network increasing the muscle activity, in concert with the left one. (B) EMG results from the left active (LA; blue), right passive (RP; green), and in-phase (IN; red) tasks. Small bars on the graph indicate standard errors of the mean across participants. *: p < 0.005, **: p < 0.001.

Recently, an intervention to restore hand motor function in hemiplegic patients by electrically contracting the muscles of the paralyzed hand in accordance with the movement of the intact hand (= contralaterally controlled functional electrical stimulation [FES]) has been shown to be effective (Knutson et al., 2007, 2012, 2022; Cunningham et al., 2019; Huang et al. al., 2021; Loh et al., 2022; Halawani et al., 2024). One may assume that the essence in this intervention is proprioceptive intervention of a hand synchronized with voluntary movement of the other hand (bilateral proprioceptive-motor coupling). However, the underlying neuronal substrates for this success has not been well described. Therefore, we first show in healthy volunteers that passive movement of one hand synchronized with voluntary movement of the other hand can effectively induce muscle activity in the former, and its underlying neural substrates in the bilateral motor cortices.

When a healthy person voluntarily moves the left hand, the right motor cortex is activated and the left motor cortex is deactivate (inhibited) due to interhemispheric inhibition. Similarly, when the right hand is moved passively, a proprioceptive signal activates the left motor cortex via somatosensory area 3a and the cerebellar vermis (Naito et al., 2016), leading to inhibition of the right motor cortex through interhemispheric inhibition (Naito et al., 2021).

We hypothesize that simultaneous voluntary movement of the left hand and passive movement of the right hand will lead the left and right motor cortices to a bilaterally active mode (Figure 1A). In this situation, interhemispheric functional connectivity must be enhanced between the motor cortices. This enhanced functional coupling may allow the bilateral motor cortices to share the voluntary motor signal input to the right motor cortex and the proprioceptive input to the left motor cortex, promoting sensory-motor associations between them. If the sensory-motor association occurs in the left motor cortex, the voluntary motor signal may effectively activate the intrinsic transcortical circuit between the left motor cortex and the right hand muscles, for instance (Fetz et al., 1980; Cheney and Fetz, 1984), increasing muscle activity during passive movement. If this association also occurs in the right motor cortex, the cortex might become part of the network increasing muscle activity, in concert with the left one.

### 2.1 Methods and results

We tested these hypotheses in healthy adults. The details of methods are described in following Supplementary Material. We recruited 55 healthy right-handed young adults (37 male, 18 female; age 19–26 years old). Their handedness was assessed by the Edinburgh Handedness Inventory (Oldfield, 1971). The motor tasks consisted of (1) a left finger active extension task (left active; LA) in which the blindfolded participants extended their left index finger to a 1 Hz tone, (2) a right finger passive extension task (right passive; RP) in which the experimenter extended the right relaxed index finger to a 1 Hz tone, and (3) an in-phase task (in-phase; IN) in which the experimenter extended the right relaxed finger (RP) synchronized with the participant’s active left finger extension (LA; Figure 1B). The study protocol was approved by the Ethics Committee of the National Institute of Information and Communications Technology, and the MRI Safety Committee of the Center for Information and Neural Networks (CiNet; no. 2003260010). We explained the details of the present study to all participants before the experiment, and they then provided written informed consent. The study was conducted according to the principles and guidelines of the Declaration of Helsinki (1975).

We first examined if muscle activity in the right relaxed finger increases during the IN task. In all tasks, surface electromyograms (EMGs) were recorded from the finger extensor muscles of the right hand. The 20-s task was repeated eight times with a 10-s rest phase in between. The root-mean-square EMG values from the first 2–19 s during the task and from the first 2–7 s during the rest phase were calculated, and average values for the eight tasks and rest phases were calculated for each individual. A total of 44 of all participants showed EMG increase during the task phase compared to the rest phase in the IN task. A two-factorial analysis of variance (repeated measurement) for tasks (3) x period (2: task-rest) revealed a significant interaction (F(2, 108) = 9.21; p < 0.001). A post hoc test revealed a significant increase in muscle activity during the IN task compared to the rest phase (p < 0.001 Bonferroni corrected); further, activity during the IN task was significantly higher than during the LA and RP tasks (p < 0.001, p < 0.005 Bonferroni corrected, respectively). In other words, the IN task could effectively increase muscle activity in the right relaxed finger, an effect that could not be induced by passive movements alone (Figure 1B).

To investigate the neural substrates underlying the IN task, brain activity was measured with a 3 Tesla functional MRI during the above three tasks. Using an image analysis software of statistical parametric mapping, we preprocessed individual images, spatially smoothed them with a 4 mm Gaussian filter, and conducted statistical analyses (see Supplementary Material). We adopted a voxel-wise threshold of p < 0.005, and evaluated the significance of brain activations in terms of the spatial extent of the activations in the entire brain or in small volume correction (SVC; p < 0.05, family-wise-error (FWE) corrected).

When comparing the IN task with the LA and RP tasks, bilateral motor-cortical activations were observed (Figure 2A red). The peaks of these activations were located in M1 (cytoarchitectonic areas 4a/4p; MNI coordinates x, y, z = −38, −16, 50 and 36, −16, 52). Specifically, in the LA task, the right M1 was activated and the left M1 was deactivated while, in the RP task, the opposite was observed (Figure 2B). Hence, when performing the LA or RP task alone, there is deactivation by interhemispheric inhibition, but the IN task, which combines LA and RP tasks, bilaterally activates motor cortices (Figure 2A, B). Since these activations were observed only during the IN task, these can be considered IN task-related motor-cortical activations, and likely associated with sensory-motor association.

**Figure 2.**
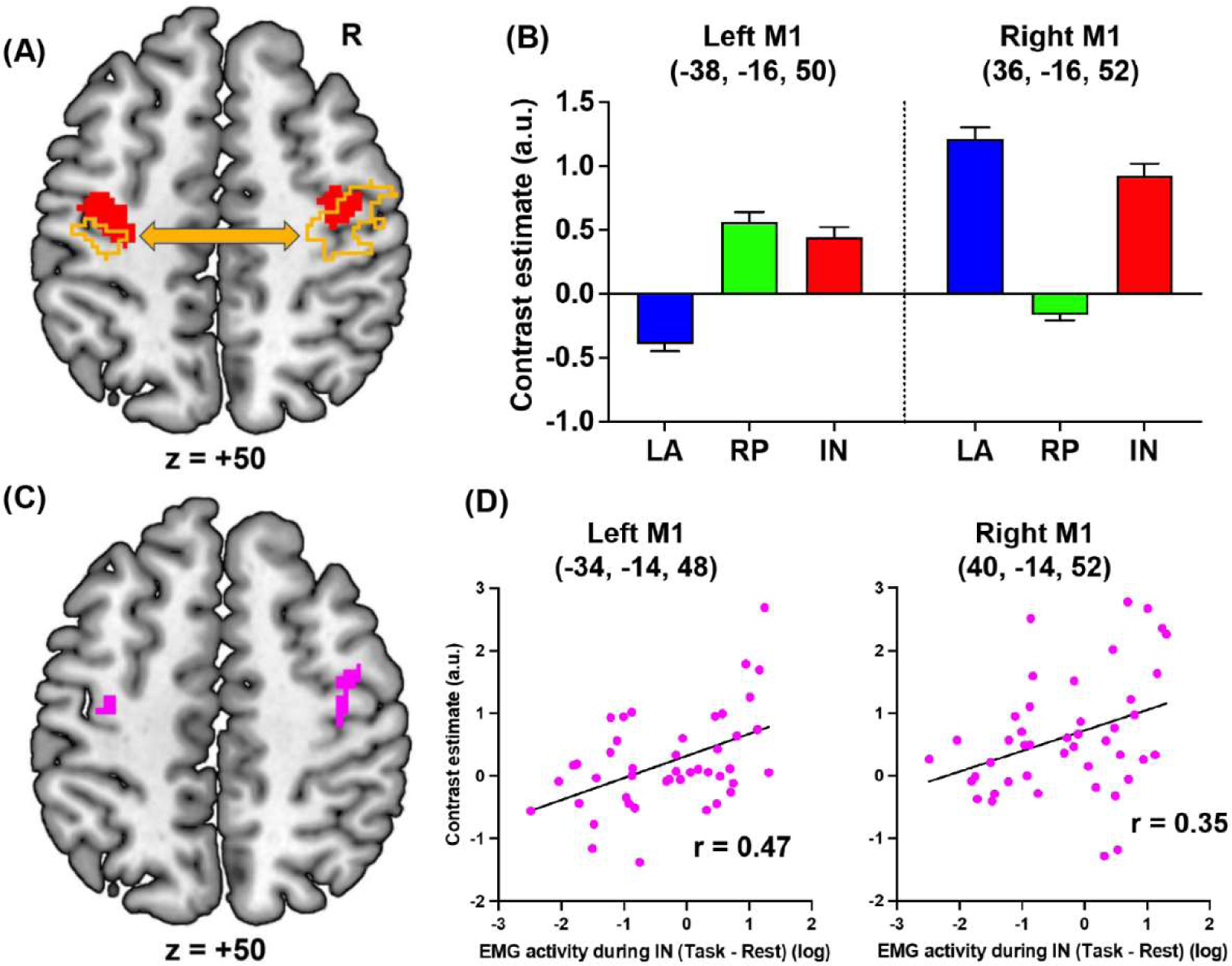
fMRI results. (A) Bilateral motor-cortical activations (red) when comparing the IN task with the LA and RP tasks. The left and right motor cortices (orange) in which activity enhanced functional coupling with the right or left motor-cortical cluster (red) respectively. (B) Brain activity in the peaks of the left and right motor-cortical clusters in each task. (C) The left and right motor cortices (pink) in which activity significantly correlated with the EMG activity during the IN task. (D) Interparticipant correlation between the left or right M1 activity and log value of EMG activity. Abbreviations: LA, left active; RP, right passive; IN, in-phase; M1, primary motor cortex; R, right; a.u., arbitrary unit.

Next, using a generalized psychophysiological interaction analysis (McLaren et al., 2012), we further examined the brain regions where activity enhanced functional coupling with the activity in the left or right motor-cortical cluster (Figure 2A red) during the IN task when compared to the other tasks (see Supplementary Material). The right (2202 voxels; peak coordinates = 34, −18, 50, area 4p) or left (746 voxels; peak coordinates = −38, −20, 52, area 4p) motor-cortical activity increased their functional coupling with the left or right cluster respectively (Figure 2A orange).

Finally, we examined if the IN task-related motor-cortical activity (IN > LA + RP) correlates with the EMG increase (task > rest) in the right extensor muscles during the IN task across the 44 participants (see Supplementary Material). In the left motor cortex, we found a significant cluster of 28 voxels (peak coordinates = −34, −14, 48; Figure 2C left pink section), which became significant after SVC (p = 0.021 FWE-corrected) within a sphere with 8 mm radius around the peak of the IN task-related left M1 activity. Plotting this relationship across participants showed that the IN task-related left M1 activity was greater in participants with higher EMG activity during the IN task (r = 0.47; Figure 2D left panel). Similarly, in the right motor cortex, we also found a significant cluster of 27 voxels (peak coordinates = 40, −14, 52; Figure 2C right pink section), which became significant after SVC (p = 0.023 FWE-corrected) within a sphere with 8 mm radius around the peak of the IN task-related right M1 activity. The IN task-related right M1 activity was also greater in participants with higher EMG activity during the IN task (r = 0.35; Figure 2D right panel). The results indicate that the IN task-related bilateral motor-cortical activities are related to the EMG increase during the IN task.

### 2.2 Discussion and possible clinical application

The present study has clearly demonstrated in healthy volunteers that passive movement of a relaxed hand synchronized with voluntary movement of the other hand can effectively induce muscle activity in the former. This intervention leads the motor cortices to bilaterally active mode, and enhances their interhemispheric functional coupling. Further research is needed to determine how the putative sensory-motor association leads to the EMG increase. However, in the current work, not only the left (contralateral) but the right (ipsilateral) motor-cortical activity correlated with the EMG increase of the right relaxed hand (Figure 2C, D). The causal relationship between these activities and the EMG increase through descending pathways remains unclear. In primates, neurons in the ipsilateral motor cortex projects to spinal interneurons (Morecraft et al., 2013, 2019). Therefore, not only the contralateral but the ipsilateral activity may be directly involved in the EMG increase through the ipsilateral descending pathway. If so, the IN task could promote activity in this pathway, which could compensate hand motor function after contralateral stroke (Lotze et al., 2006).

The current results were obtained in healthy young participants; thus, the current intervention needs to be tested in stroke patients. There are several caveats to be considered when applying our intervention clinically. First, the patients must have intact proprioceptive pathways on the paralyzed side, since proprioceptive input from the paretic hand (most likely synchronized with the movement of the intact hand) could be crucial. However, even when proprioceptive pathways are damaged, viewing the movement of one’s own paretic hand synchronized with the voluntary movement of the intact hand might cause similar effects to bilateral mirror therapy (Thieme et al., 2018; Bello et al., 2020; Zhang et al., 2024). Second, complete damage to the motor cortex severely impairs both motor and proprioceptive processing (Naito et al., 2011). In this study, we confirmed that a right-handed 71-year-old male patient with a focal subcortical hemorrhage over the left precentral hand region was unable to move his right arm/hand/fingers and could not experience proprioceptive illusory movement of the right hand on the third week after the stroke (These functions were improved six months after stroke). On the other hand, in the case of partial motor-cortical damage by an experimental ischemic block, the motor and proprioceptive functions seems to be compensated by spared adjacent tissue around stroke core (Nudo, 2003). Hence, in the latter case, capability of residual tissue associating motor and proprioceptive signals would be the key.

Finally, we expect that the IN task may be effective in the acute and subacute phases of stroke when the brain spontaneously shifts to the bilaterally active mode (Grefkes and Fink, 2020). We assume that moving the immobile hand from the acute and subacute phases of stroke may decrease the risk of excessive interhemispheric inhibition from the contralesional (intact) motor cortex to the ipsilesional one (Grefkes and Fink, 2020). In addition, such early phase intervention might decrease the risk of spasticity or rigidity progress caused by long-term immobility of the paretic hand.

Recently, we have developed a new intervention using a haptic robotic system, which utilizes the plastic capability in the interhemispheric interaction shown in the above bilateral proprioceptive-motor coupling intervention. This system can measure the movements of individual fingers of one hand online, and simultaneously (delay about 20 ms) copy these movements to the corresponding fingers of the other hand, automatically moving them. We are now evaluating its effectiveness for restoration of hand/finger motor functions, and its underlying neural substrates. This system is easy to wear and to be prepared, and can be utilized even from the acute phase (even at bed side). In this intervention, we ask stroke patients to try voluntarily moving both intact and paretic (immobile) hands simultaneously, and the paretic hand movement is assisted by their own voluntary movement of the intact hand. Therefore, stroke patients can directly experience and see movements of their paretic hand synchronized with voluntary movements of their intact hand. We expect that this intervention generates better outcome than mirror-induced visual illusion therapy, and has higher convenience in preparation and less uncomfortable (painful) effect compared to the above contralaterally controlled FES. This intervention seems to be a very promising so far, by facilitating voluntary movement of the paretic hand after the intervention. We will share the results in our near future.

## 3 Improvement of hand dexterity utilizing left-right independent mode

The bilaterally active mode is accompanied by involuntary muscle activity and movement. After stroke, when the motor cortex ipsilateral to the hand is active, involuntary mirror movements and muscle activity of the opposite hand can occur (Jang et al., 2004; Carson, 2005). Involuntary mirror movements and muscle activity can also be observed in children with immature interhemispheric inhibition between the left and right motor cortices and in older adults with reduced interhemispheric inhibition (Addamo et al., 2013). Such involuntary movements are more likely to appear when the bilaterally active mode is overtrained. In addition, excessive activity in the motor cortex ipsilateral to the hand in children and older adults is closely related to their lower hand dexterity (Naito et al., 2020, 2021). To circumvent this, training the motor cortex into a left-right independent mode, by training the left and right hands to perform different movements, may be effective.

In our previous 2-month intervention study in healthy older adults (Naito et al., 2021), we examined if the bimanually different movement training can reactivate motor-cortical interhemispheric inhibition declined with age in a way that reduces excessive activity of the ipsilateral motor cortex during a non-motor proprioceptive task of the right hand, and if such activity reduction is related to dexterity improvement of the hand. The older adults (over 65 years old) were divided into two groups: one group was a bimanual group in which the left and right hands were trained to perform different movements (including left-right out-of-phase movements) every day, and the other group was a right-handed group where the right hand alone was trained to perform the movements of the bimanual group. The dexterity of the right fingers was assessed using the 12-hole peg test.

Although none of the groups trained on the peg test per se during the 2-month period, only the bimanual group showed significant improvement in peg task performance (= dexterity) after training. Further, only this group showed significant reduction of excessive ipsilateral motor-cortical activity during the proprioceptive task (most likely due to reactivation of interhemispheric inhibition from the left motor cortex to the right one) after the training. This effect was more pronounced in the older adults, whose right motor cortex was overactive (= loss of inhibition from the left motor cortex) before the training, and the degree of reduction in the right motor-cortical activity was significantly correlated with the dexterity improvement of the right hand.

The bilaterally active mode of the left and right motor cortices observed in healthy older adults is similar to that observed after stroke. This suggests that stroke patients who can move their paretic hands to some extent can improve their hand dexterity by actively training the left and right hand to perform different movements.

## 4 Conclusion

Our recent EMG and functional MRI study in healthy younger adults suggests that proprioceptive intervention of the paralyzed (immobile) hand synchronized with voluntary movement of the intact hand (bilateral proprioceptive-motor coupling intervention) may facilitate restoring motor function of the paretic hand after stroke, utilizing the spontaneous bilaterally active mode of motor cortices frequently observed after stroke (Figure 3). This way, muscle activity could be induced in the paretic hand through the association between voluntary motor signals and proprioceptive inputs to the motor cortex. In addition, our previous functional MRI study in healthy older adults indicates the possibility that, when a stroke patient becomes able to manage to move the paretic hand but the movement is clumsy, training both hands to perform different movements (bilaterally different movement training) that guides the motor cortices into left-right independent mode could improve interhemispheric inhibition and hand dexterity (Figure 3).

**Figure 3.**
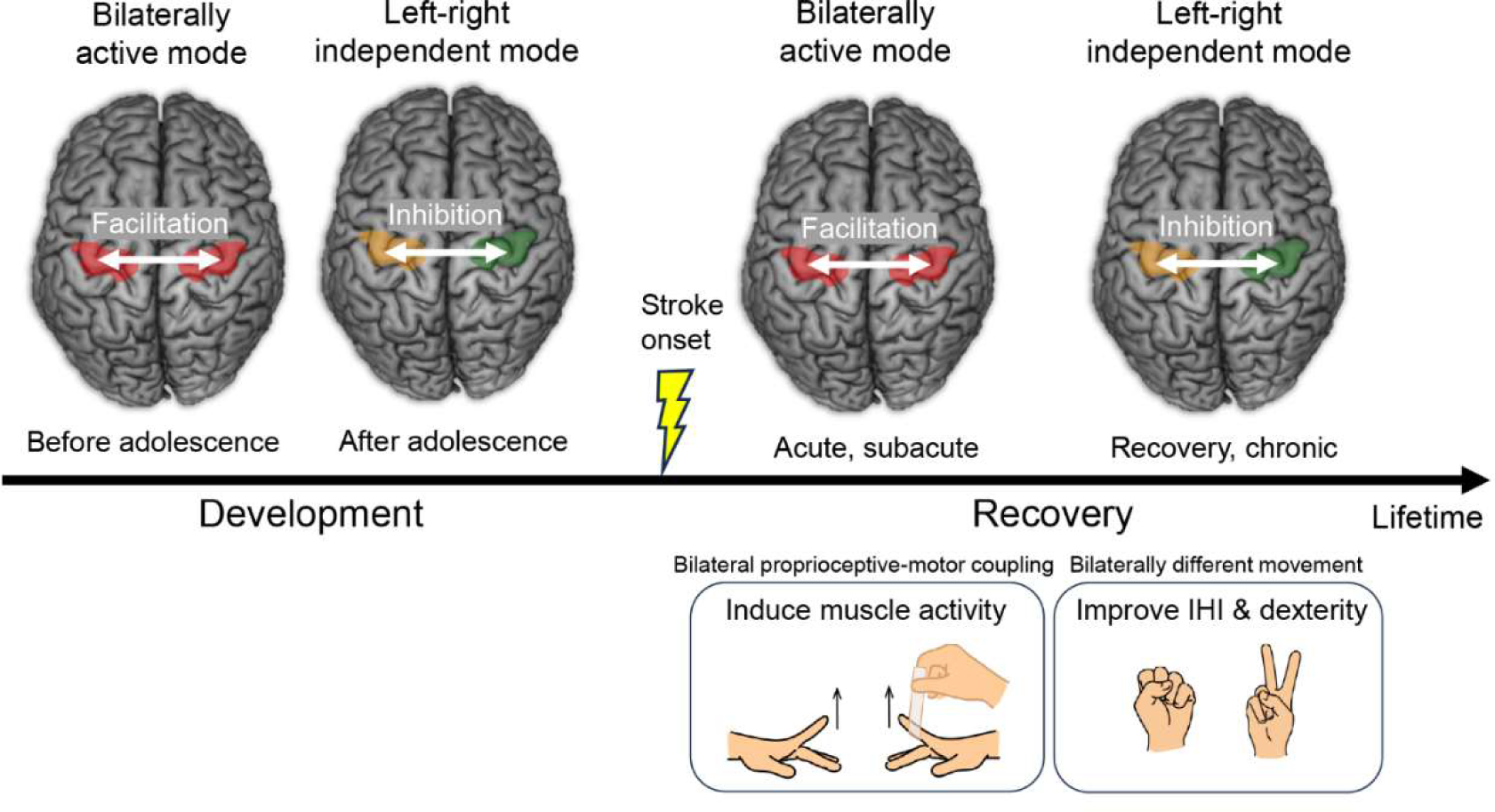
Transition of the left and right motor cortices from the bilaterally active mode to the left-right independent mode. This is a rule of typical development of hand motor function and also a common feature of motor function recovery after stroke. Proposed interventions during recovery are shown at right bottom. Abbreviations: IHI, interhemispheric inhibition.

The transition of the motor cortices from the bilaterally active mode to the left-right independent mode is a common feature of motor function recovery after stroke (Jang et al., 2004; Grefkes and Fink, 2020), and also is a rule of development of hand motor function (Figure 3), which everyone has experienced during the developmental process (Morita et al., 2019, 2021). The habilitation of “re-habilitation” means “to gain ability,” and it is during the developmental period that the most ability is gained in a person’s lifetime. Therefore, rehabilitation according to developmental rules may be the most natural and effective way for the brain to reacquire abilities. When planning rehabilitation strategies, taking advantage of the brain’s spontaneous recovery process and referring to its developmental rules may be important considerations.

## Conflict of Interest

The authors declare that the research was conducted in the absence of any commercial or financial relationships that could be construed as a potential conflict of interest.

## Author Contributions

All authors contributed to conceptualization, validation and writing - review editing, and investigation. HN, YT, and TM: formal analysis, HN, TM, and EN: funding acquisition. HN, YT, and EN: visualization. EN: writing - original draft preparation, project administration, and supervision. All authors contributed to the article and approved the submitted version.

## Funding

This study was supported by JSPS KAKENHI Grant Nos. JP19H05723, JP23H03706, and JP23K17453 for EN, and by JSPS KAKENHI Grant No. JP20H04492 for TM, and by JSPS KAKENHI Grant No. JP23K10417 for HN.

## Acknowledgments

The funding sources were not involved in the study design; collection, analysis, interpretation of data; writing of the report; or the decision to submit the article for publication. The authors are grateful to Dr Tsuyoshi Ikegami and Dr Jihoon Park for their valuable comments on this work. They also thank Dr Tadashi Isa and Dr Hidenori Aizawa for intensive discussion about neural substrates of disinhibition of interhemispheric inhibition through “Hyper-adaptation” project (JP19H05723), CiNet MRI staff and Ms Mayumi Irikawa for their experimental supports, and Ms Keiko Ueyama for the illustration.

## Data Availability Statement

Datasets are available on request: The raw data supporting the conclusions of this article will be made available by the corresponding author (EN).

## Supplementary methods

### Participants

A total of 55 healthy right-handed young adults (37 men, 18 women: mean age, 21.9 [mean] ± 1.6 [standard deviation]; range, 19–26 years) participated in this study. None of them had a history of self-reported neurological, psychiatric, or motor disorders. Their handedness was confirmed using the Edinburgh handedness inventory (Oldfield, 1971). The study protocol was approved by the Ethics Committee of the National Institute of Information and Communications Technology and the MRI Safety Committee of the Center for Information and Neural Networks (CiNet; no. 2003260010). Details of the experiment were explained to each participant before the experiment, and they provided written informed consent. This study was conducted in accordance with the principles and guidelines of the Declaration of Helsinki (1975).

### Motor tasks

We prepared three motor tasks (Figure 1B). In a left finger active extension (left active; LA) task, the participants with their eyes closed extended their left index finger in synchronization with a computer-generated 1 Hz tone. They were asked to repeat 1 Hz extension and its following relaxation of the left finger, while relaxing their right hand. They were encouraged to extend the left finger to its maximum angle. In a right finger passive extension task (right passive; RP), experimenters extended their right relaxed index finger in synchronization with the 1 Hz tone. The blindfolded participants were asked to relax both hands, the experimenters (HN during EMG recoding and EN during fMRI experiment) repeated 1 Hz extension of the right finger to its maximum angle. In an in-phase (IN) task, the experimenters extended the right relaxed finger (RP) at 1 Hz synchronized with the participant’s active 1 Hz left finger extension (LA). Before we started the tasks, we measured maximum extension angles of the left and right index fingers in each participant using an angle measuring device (easyangle, ITO PHYSIOTHERAPY&REHABILITATION Co., Ltd., Tokyo, Japan). The mean maximum angles across participants were 40.0 ± 8.0 and 40.4 ± 7.5 degrees for the left and right fingers, respectively.

All of the tasks were performed both during EMG recording and during fMRI experiment. The order of the task (IN, RP, and IN) was fixed across all participants, as individual difference in EMG activity was also one of our research purposes. Both in the EMG recording and in the fMRI experiment, each participant completed one experimental run for each task. In a run, 20-s task epoch was repeated eight times with a 10-s rest phase in between. Each run also included a 10-s rest phase before the start of the first task epoch. The participants and experimenters were given computer-generated auditory instructions (“3, 2, 1, start” and “stop”) to inform them of the start and finish of a task epoch.

### EMG recording and analysis

EMG recording was conducted before the fMRI experiment outside the MRI scanner room. Each participant was seated in a chair. The left and right forearms were placed on a desk with towels laid on it, with their palms down. The mean distance between two hands across participants was 32.5 ± 6.3 cm.

In each task, surface electromyogram (EMG) was recorded from the right extensor digitorum communis (EDC). The skin surface was cleaned with gel (Nuprep, Nihonsanteku Co., Ltd., Osaka, Japan). Using double-sided adhesive tape (Map-3M402, Nihonsanteku Co., Ltd., Osaka, Japan), a bipolar Ag-AgCl surface electrode (BA-U410 m(A)-015, Nihonsanteku Co., Ltd., Osaka, Japan) was placed over the belly of the muscle. The EMG signals were recorded using the EMG Multi Analysis Programe MaP1038L (Nihonsanteku Co., Ltd., Osaka, Japan; sampling rate = 2048 Hz; high frequency filter = 1000 Hz; time constant = 0.03 sec). The EMG raw data were amplified (gain, 10000), and smoothed using a moving average. The root-mean-square EMG values from the first 2–19 s during the task epoch and from the first 2–7 s during the rest phase were calculated, and average values for the eight tasks and rest phases were calculated for each participant. For statistical analysis, a two-factorial analysis of variance (repeated measurement) for tasks (3) x period (2: task-rest) was performed (Figure 1B).

### fMRI experiment

The participants were placed in the supine position in the MRI scanner, and performed three motor tasks. The head was secured with a sponge cushion and adhesive tape, and the participants wore MRI-compatible earphones (SENSIMETRICS Model S14). The arms were set in a natural semi-rotated position, stretched along the body and relaxed. Both the left and right hands were placed on hand cushions and relaxed with their palms down. They were instructed to close their eyes, relax their entire bodies without producing unnecessary movements, and think only of things relevant to the tasks assigned. We told them to close their eyes just before each experimental run.

### fMRI acquisition

Functional images were acquired using T2*-weighted gradient echo-planar imaging (EPI) sequences on a 3.0-Tesla MRI scanner (Vida; Siemens, Germany) equipped with a 64-channel array head-neck coil. A multiband imaging technique was used (multiband factor, 3; Moeller et al. 2010). Each volume consisted of 51 slices (slice thickness, 3.0 mm with no inter-slice thickness) acquired in an interleaved manner, covering the entire brain. The time interval between successive acquisitions from the same slice (TR) was 1,000 ms. An echo time (TE) of 30 ms and a flip angle (FA) of 60° were used. The field of view (FOV) was 210 x 210 mm^2^ and the matrix size was 70 x 70 pixels. The voxel dimensions were 3 × 3 × 3 mm^3^ in the x-, y-, and z-axes, respectively. Each participant completed one 250-s experimental run for each task (see above), which included a 10-s period (before the first rest phase) that was excluded from the analysis to eliminate the influence of unsteady magnetization.

Prior to EPI acquisition, filed map images were obtained using a gradient echo sequence (TR = 1000 ms, TE1 = 5.29 ms or TE2 = 7.75 ms, FOV = 210 x 210 mm^2^, FA = 60°, matrix size = 70 x 70 pixels, number of slices = 51, voxel dimensions = 3 × 3 × 3 mm^3^), in order to correct geometrical distortion of EPI images caused by static field inhomogeneities (see below).

As an anatomical reference, a T1-weighted magnetization-prepared rapid gradient echo (MP-RAGE) image was acquired using the same scanner. The imaging parameters were as follows: TR = 1900ms, TE = 2.48ms, FA = 9°, FOV = 256 × 256 mm^2^, matrix size = 256 × 256 pixels, slice thickness = 1.0 mm, voxel size = 1 × 1 × 1 mm^3^, and 208 contiguous sagittal slices.

### fMRI analysis Preprocessing

To eliminate the influence of unsteady magnetization during the tasks, the first 10 volumes (10 second) of EPI images in each run were excluded from the analysis. Acquired imaging data were analyzed using SPM 12 (default setting: Wellcome Trust Centre for Neuroimaging, London, UK) running on MATLAB R2017a (MathWorks, Sherborn, MA, USA).

A voxel-displacement map (VDM) was calculated from the acquired field map images. EPI images were aligned to the first image (realigned images). Time series data of the head position during the fMRI experiment were obtained by a rigid body transformation (linear transformation) using the least squares method for six realign parameters (translation along the x-, y-, and z-axes and the rotational displacements of pitch, raw, and roll). Then, the head movements of the participants were evaluated by the framewise displacement (FD) values based on the six parameters (Power et al., 2012). To inspect FD values through all frames of an entire experimental run, we counted the number of frames that had an FD of over 0.9 mm in each participant according to a previous study (Siegel et al., 2014). None of the participants showed FD values that exceeded 0.9 mm in more than 5 % of the total volumes, and thus none of them were excluded from the following analyses.

The realigned images were unwarped using the VDM to correct geometrical distortion of these images. In each participant, the T1-weighted structural image was co-registered to the mean image of all realigned and unwarped EPI images. The individual co-registered T1-weighted structural image was spatially normalized to the standard stereotactic Montreal Neurological Institute (MNI) space (Evans et al., 1994). Applying the parameter estimated in this process, the individual realigned and unwarped images were normalized to the MNI space with 2-mm isotropic voxel size using the SPM12 normalization algorithm. Finally, the normalized images were filtered using a Gaussian kernel with a full-width at half-maximum (FWHM) of 4 mm along the x-, y-, and z-axes.

### Single-subject analysis

After preprocessing, we first explored task-related activations and deactivations in each participant with a general linear model (Friston et al., 1995; Worsley & Friston, 1995). This was done for each task. For the first-level analysis, a design matrix was prepared for each participant. The design matrix contained a boxcar function for the task epoch that was convolved with a canonical hemodynamic response function (HRF). Six realignment parameters were also included in the design matrix as regressors to correct for residual motion-related noise after the realignment. Contrast images showing activation (task > rest) and deactivation (rest > task) in each task and the image of IN vs. LA + RP were created for each participant, which was used in the following second-level group analysis. In the first-level analysis, global mean scaling was not performed to avoid inducing type I error in the assessment of negative blood oxygenation-level dependent (BOLD) responses (Aguirre et al., 1998).

### Group analysis

In the second-level group analysis, we first examined activations and deactivations in each task (not shown in Figure). Next, we examined if the bilateral motor cortices are active during the IN task when compared with the LA and RP tasks (IN > LA + RP). In these analyses, we generated a voxel-cluster image using an uncorrected voxel-wise threshold of p < 0.005, and evaluated the significance of brain activations in terms of the spatial extent of the activations in the entire brain (p < 0.05, family-wise-error (FWE) corrected). Since our main regions-of-interest were the bilateral motor cortices (dorsal premotor cortex and primary motor cortex), the present study only reported motor-cortical activations (Figure 2A red sections). For anatomical identification of the activation peaks, we referred to the cytoarchitectonic probability maps (Eickhoff et al., 2005; Amunts et al., 2020).

In order to visualize task-dependent activity change in the identified bilateral motor-cortical clusters (Figure 2A red sections), we extracted individual brain activity (parameter estimates) from the peaks of left (x, y, z = −38, −16, 50) and right (36, −16, 52) motor-cortical clusters for each task, and calculated the mean across participants (Figure 2B).

### Task-related functional connectivity analysis

We further examined the brain regions where activity enhanced functional coupling with the activity in the left or right motor-cortical cluster (seed regions; Figure 2A red) during the IN task when compared to the other (LA and RP) tasks, by conducting a generalized psychophysiological interaction analysis (gPPI) (McLaren et al., 2012). Defining seed regions from the contrast analysis and using the seed regions in the functional connectivity analysis is completely different from the circular analysis where a ROI is defined by an identified cluster in one contrast of activation analysis and then used for another contrast in the same analysis (Wei et al., 2022). This analysis was performed on SPM-preprocessed fMRI data using the CONN toolbox version 20.b (Whitfield-Gabrieli & Nieto-Castanon, 2012; Nieto-Castanon, 2020). Physiological noises originating from the white matter and cerebrospinal fluid were removed using the component-based noise correction method (CompCor) in the toolbox (Behzadi et al., 2007). Head motion-related artifacts, scrubbing, and condition effects were also removed. A temporal band-pass filter of 0.008–0.09 Hz was applied, because we wanted to examine task-related functional connectivity change in this slower range of brain activity fluctuation below than the cardiac and respiratory cycles (0.1–1.2 Hz) (Cordes et al. 2001).

In the gPPI analysis, we used each of the left and right motor-cortical clusters as a seed region. In each participant, the time course of the average fMRI signal across the voxels in each seed region was deconvolved using the canonical HRF (physiological variable). Then, we performed a general linear model analysis using the design matrix and included the following regressors: physiological variable, boxcar function for the task epoch (psychological variable), and multiplication of the physiological variable and the psychological variable (PPI). These variables were convolved with a canonical HRF. Six realignment parameters were also included in the design matrix as regressors of no interest.

In each task, we first generated an image of voxels showing to what extent their activities changed with the PPI regressor of each seed region in each participant. Then, we generated a contrast image (IN > LA + RP) that shows the IN task-related connectivity change for each participant. We used this individual image in the second-level group analysis. In the second-level analysis, we searched for significant clusters in the entire brain (uncorrected voxel-wise threshold of p < 0.005 and extent threshold of p < 0.05 FWE-corrected). We only reported the clusters identified around the bilateral motor cortices (Figure 2A orange) in this study.

### Correlation analysis

Finally, we examined if the IN task-related motor-cortical activity (IN > LA + RP) correlates with the EMG increase (task > rest) in the right extensor muscles during the IN task across participants. A correlation analysis was conducted using individual EMG activity measured outside MRI as a covariate. Since 44 of 55 participants showed EMG increase during the task epoch compared to the rest phase during the IN task, the correlation analysis was done for the 44 participants. The individual raw data of EMG increase were not normally distributed, therefore, the logarithm of the data was calculated and used as covariates. We first generated a cluster image using uncorrected voxel-wise threshold of p < 0.005. Since our main interest was the bilateral motor cortices, in the statistical evaluation, we conducted small volume correction (SVC; p < 0.05 FWE-corrected) approach, using a sphere with 8 mm radius around the peak (−38, −16, 50) of the IN task-related left M1 activity or a sphere with 8 mm radius around the peak (36, - 16, 52) of the IN task-related right M1 activity respectively (Figure 2C). The 8 mm radius was selected based on final smoothness of functional image (maximum 7.1 mm FWHM). We extracted individual brain activity (parameter estimates) from a peak of each identified cluster (−34, −14, 48 for the left and 40, −14, 52 for the right), and displayed interparticipant correlation between the brain activity and the logarithm of EMG activity (Figure 2D).

## Notes

### Competing Interest Statement

The authors have declared no competing interest.

